## Supplemental for "Intra and interspecific diversity in a tropical plant clade alter herbivory and ecosystem resilience"

**Supplementary figures and tables**

**Table S1.** *Study site characteristics and experimental details*

|  | La Selva Biological Station | Yanayacu Biological Station | El Fundo Génova | Mogi-Guaçu Biological Reserve | Uaimii  State Forest |
| --- | --- | --- | --- | --- | --- |
| Location | N9.9400, W84.0433 | S0.6008, W77.8903 | S11.0948, W75.3516 | S32.2510, W47.1558 | S30.2957,  W43.5731 |
| Mean elevation (m a.s.l.) | 35 | 2124 | 1133 | 627 | 800 |
| Mean annual precipitation (mm) | 4495 | 2900 | 1767 | 1271 | 1600 |
| Mean annual T (°C) | 25 | 16 | 23 | 26 | 19 |
| Date planted | 2015-03-01 | 2015-07-01 | 2015-10-01 | 2017-02-01 | 2018-02-20 |
| Date harvested | 2017-02-01 | 2016-12-01 | 2017-07-01 | 2019-12-01 | 2019-12-20 |
| Number of species in high richness plots | 12 | 12 | 6 | 4 | 3 |
| Planted in pots / ground | Pots | Pots | Pots | Ground | Ground |
| Water addition treatment applied | Yes | Yes | Yes | No | No |
| Number of *Piper* planted | 360 | 360 | 360 | 432 | 144 |
| Number of *Piper* surviving till the end of the experiment | 200 | 259 | 56 | 182 | 123 |

*Climate data from La Selva Biological station are for the experimental period (2015-2018) and were provided by the Organization for Tropical Studies. Data from El Fundo Génova are based on nearby San Ramón (<https://en.climate-data.org/south-america/peru/junin/san-ramon-28556/>). Data from Ecuador were provided by Yanayacu Biological Station. Data for Mogi-Guaçu Biological Reserve are from January 2017-December 2019 and are from the Centro Integrado de Informações Agrometeorológicas of São Paulo. Data for Uaimii State Forest are from the Plano de Manejo FLOE Uaimii, Instituto Estadual de Florestas of Minas Gerais

**Table S2.** Species of *Piper* used at each study location

| Study location | | | | |
| --- | --- | --- | --- | --- |
| La Selva Biological Station | Yanayacu Biological Station | El Fundo Génova | Mogi-Guaçu Biological Reserve | Uaimii  State Forest |
| arboreum | grande | arboreum | arboreum | lepturum |
| peltatum | peludo | reticulatum | crassinervium | corcovadense |
| biolleyi | ecuadorense | glabribaccum | richardifolium | vicosanum |
| sancti-felicis | perareolatum | chanchamayanum | miquelianum |  |
| reticulatum | baezanum | armatum |  |  |
| nudifolium | escabrosa | lechlerianum |  |  |
| multiplinervium | kelleii |  |  |  |
| decurrens | hispidum |  |  |  |
| pseudobumbratum | schupii |  |  |  |
| imperiale | pubinervulum |  |  |  |
| garagaranum | pequenia |  |  |  |
| urostachyum |  |  |  |  |

**Table S3.** Mean parameter estimates and probability of direction (PD) for the effects of increases in intraspecific diversity, interspecific richness, water availability and insect richness on measures of herbivory, plant mortality, and insect richness

| Site | Predictor variable | Response variable | Mean parameter estimate | PD |
| --- | --- | --- | --- | --- |
| All | Intraspecific richness | Percent herbivory | -0.87 | 63.59% |
| All | Intraspecific richness | Presence of damage | -3.26 | 71.60% |
| All | Intraspecific richness | Variance in herbivory | -9.97 | 57.31% |
| All | Intraspecific richness | Insect richness | -0.02 | 59.41% |
| All | Intraspecific richness | Plant survival | 6.50 | 92.48% |
| All | Insect richness | Percent herbivory | 8.78 | 100.00% |
| All | Insect richness | Presence of damage | 6.63 | 96.73% |
| All | Insect richness | Variance in herbivory | 17.86 | 71.13% |
| All | Insect richness | Plant survival | 3.35 | 66.21% |
| All | Interspecific richness | Percent herbivory | 0.02 | 50.34% |
| All | Interspecific richness | Presence of damage | 7.40 | 83.01% |
| All | Interspecific richness | Variance in herbivory | -52.43 | 73.46% |
| All | Interspecific richness | Insect richness | 0.20 | 95.00% |
| All | Interspecific richness | Plant survival | 10.66 | 80.22% |
| All | Water availability | Percent herbivory | -4.19 | 98.73% |
| All | Water availability | Presence of damage | -6.33 | 89.46% |
| All | Water availability | Variance in herbivory | -118.10 | 91.35% |
| All | Water availability | Insect richness | -0.07 | 64.52% |
| All | Water availability | Plant survival | 5.15 | 79.29% |
| Costa Rica | Intraspecific richness | Percent herbivory | -0.10 | 52.68% |
| Costa Rica | Intraspecific richness | Presence of damage | 2.86 | 79.84% |
| Costa Rica | Intraspecific richness | Variance in herbivory | 37.95 | 72.01% |
| Costa Rica | Intraspecific richness | Insect richness | 0.10 | 98.07% |
| Costa Rica | Intraspecific richness | Plant survival | 3.72 | 80.91% |
| Costa Rica | Insect richness | Percent herbivory | 7.53 | 100.00% |
| Costa Rica | Insect richness | Presence of damage | 5.60 | 100.00% |
| Costa Rica | Insect richness | Variance in herbivory | -6.74 | 65.69% |
| Costa Rica | Insect richness | Plant survival | 3.60 | 66.25% |
| Costa Rica | Interspecific richness | Percent herbivory | -2.43 | 91.53% |
| Costa Rica | Interspecific richness | Presence of damage | -4.89 | 89.55% |
| Costa Rica | Interspecific richness | Variance in herbivory | -89.88 | 83.97% |
| Costa Rica | Interspecific richness | Insect richness | -0.08 | 93.43% |
| Costa Rica | Interspecific richness | Plant survival | 18.69 | 99.61% |
| Costa Rica | Water availability | Percent herbivory | -4.76 | 99.99% |
| Costa Rica | Water availability | Presence of damage | -7.96 | 99.90% |
| Costa Rica | Water availability | Variance in herbivory | -151.05 | 98.04% |
| Costa Rica | Water availability | Insect richness | -0.02 | 62.33% |
| Costa Rica | Water availability | Plant survival | 12.09 | 99.74% |
| Ecuador | Intraspecific richness | Percent herbivory | -3.50 | 99.87% |
| Ecuador | Intraspecific richness | Presence of damage | -8.10 | 99.95% |
| Ecuador | Intraspecific richness | Variance in herbivory | -80.24 | 92.45% |
| Ecuador | Intraspecific richness | Insect richness | -0.18 | 100.00% |
| Ecuador | Intraspecific richness | Plant survival | 20.46 | 100.00% |
| Ecuador | Insect richness | Percent herbivory | 8.68 | 100.00% |
| Ecuador | Insect richness | Presence of damage | 8.23 | 100.00% |
| Ecuador | Insect richness | Variance in herbivory | 22.77 | 97.99% |
| Ecuador | Insect richness | Plant survival | 7.46 | 81.29% |
| Ecuador | Interspecific richness | Percent herbivory | -1.74 | 89.56% |
| Ecuador | Interspecific richness | Presence of damage | 8.46 | 99.73% |
| Ecuador | Interspecific richness | Variance in herbivory | 16.01 | 57.91% |
| Ecuador | Interspecific richness | Insect richness | 0.58 | 100.00% |
| Ecuador | Interspecific richness | Plant survival | 7.78 | 85.86% |
| Ecuador | Water availability | Percent herbivory | -2.67 | 99.52% |
| Ecuador | Water availability | Presence of damage | -5.72 | 99.78% |
| Ecuador | Water availability | Variance in herbivory | -28.52 | 68.94% |
| Ecuador | Water availability | Insect richness | 0.04 | 85.57% |
| Ecuador | Water availability | Plant survival | 5.51 | 90.57% |
| Mogi | Intraspecific richness | Percent herbivory | -2.15 | 98.59% |
| Mogi | Intraspecific richness | Presence of damage | -4.29 | 97.74% |
| Mogi | Intraspecific richness | Variance in herbivory | -24.33 | 67.12% |
| Mogi | Intraspecific richness | Insect richness | 0.02 | 75.41% |
| Mogi | Intraspecific richness | Plant survival | 1.57 | 66.65% |
| Mogi | Insect richness | Percent herbivory | 11.17 | 100.00% |
| Mogi | Insect richness | Presence of damage | 12.22 | 100.00% |
| Mogi | Insect richness | Variance in herbivory | 80.94 | 99.99% |
| Mogi | Insect richness | Plant survival | 1.95 | 58.03% |
| Mogi | Interspecific richness | Percent herbivory | 2.02 | 90.42% |
| Mogi | Interspecific richness | Presence of damage | 6.39 | 97.05% |
| Mogi | Interspecific richness | Variance in herbivory | -116.52 | 90.64% |
| Mogi | Interspecific richness | Insect richness | 0.04 | 80.94% |
| Mogi | Interspecific richness | Plant survival | 17.14 | 99.05% |
| Peru | Intraspecific richness | Percent herbivory | -0.09 | 51.56% |
| Peru | Intraspecific richness | Presence of damage | -9.11 | 94.39% |
| Peru | Intraspecific richness | Variance in herbivory | -58.63 | 73.32% |
| Peru | Intraspecific richness | Insect richness | -0.03 | 63.89% |
| Peru | Intraspecific richness | Plant survival | 2.84 | 74.08% |
| Peru | Insect richness | Percent herbivory | 8.09 | 100.00% |
| Peru | Insect richness | Presence of damage | 4.38 | 99.51% |
| Peru | Insect richness | Variance in herbivory | 30.07 | 87.18% |
| Peru | Insect richness | Plant survival | -2.31 | 61.10% |
| Peru | Interspecific richness | Percent herbivory | 0.14 | 50.60% |
| Peru | Interspecific richness | Presence of damage | 16.49 | 97.23% |
| Peru | Interspecific richness | Variance in herbivory | -69.46 | 70.67% |
| Peru | Interspecific richness | Insect richness | 0.16 | 91.04% |
| Peru | Interspecific richness | Plant survival | 5.70 | 77.33% |
| Peru | Water availability | Percent herbivory | -5.15 | 99.75% |
| Peru | Water availability | Presence of damage | -5.27 | 89.79% |
| Peru | Water availability | Variance in herbivory | -173.87 | 94.77% |
| Peru | Water availability | Insect richness | -0.22 | 99.69% |
| Peru | Water availability | Plant survival | -2.10 | 69.02% |
| Uaimii | Intraspecific richness | Percent herbivory | 1.51 | 71.49% |
| Uaimii | Intraspecific richness | Presence of damage | 2.21 | 64.77% |
| Uaimii | Intraspecific richness | Variance in herbivory | 74.93 | 79.39% |
| Uaimii | Intraspecific richness | Insect richness | -0.01 | 55.41% |
| Uaimii | Intraspecific richness | Plant survival | 3.92 | 77.51% |
| Uaimii | Insect richness | Percent herbivory | 8.41 | 100.00% |
| Uaimii | Insect richness | Presence of damage | 2.83 | 85.39% |
| Uaimii | Insect richness | Variance in herbivory | -36.56 | 83.52% |
| Uaimii | Insect richness | Plant survival | 5.98 | 75.15% |
| Uaimii | Interspecific richness | Percent herbivory | 2.23 | 70.86% |
| Uaimii | Interspecific richness | Presence of damage | 10.61 | 87.21% |
| Uaimii | Interspecific richness | Variance in herbivory | -3.20 | 52.59% |
| Uaimii | Interspecific richness | Insect richness | 0.27 | 98.14% |
| Uaimii | Interspecific richness | Plant survival | 3.53 | 62.35% |

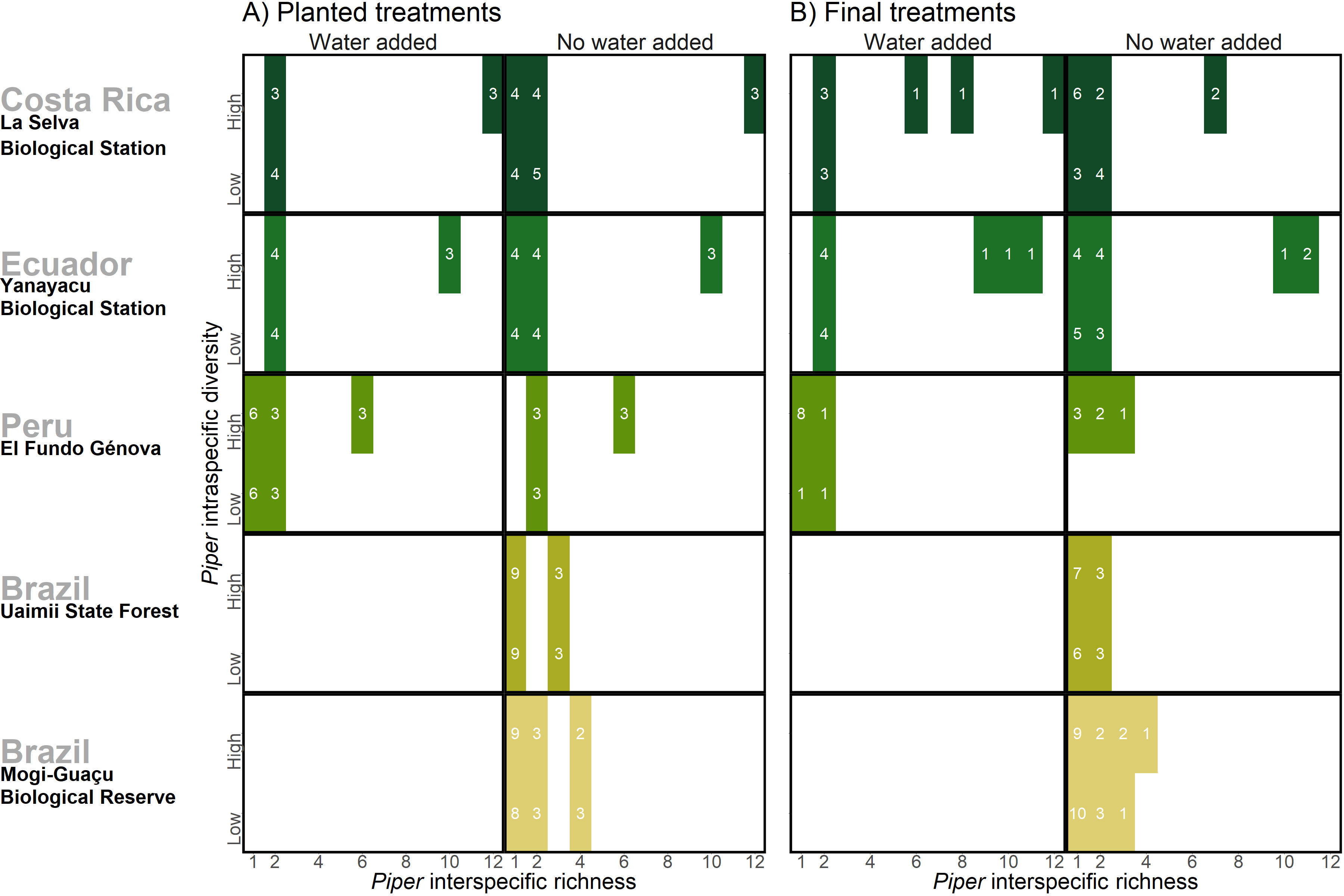
**Figure S1.** *Treatments and number of plots used across sites*

A) Treatments of intraspecific richness, interspecific richness, and water addition at the beginning of the experimental period at each of the five study sites, and B) final treatments at the end of the experimental period in each site. White numerals indicate number of plots used. Changes in *Piper* species richness and the loss of some treatment combinations was due to *Piper* mortality over the course of the experimental period

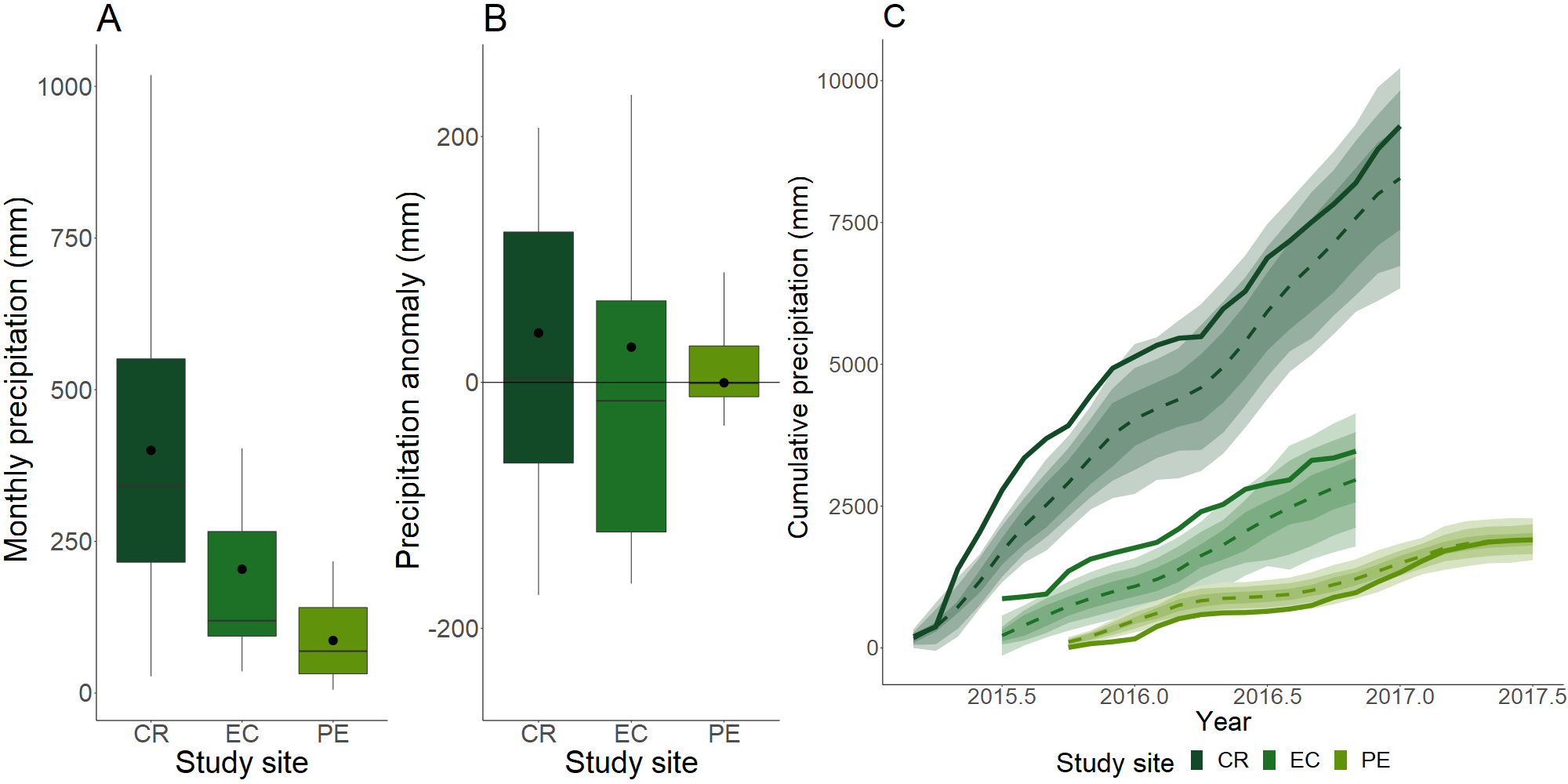
**Figure S2.** *Precipitation levels at study sites where the water addition treatment was applied*

A) Monthly precipitation over the experimental period at Costa Rica (CR), Ecuador (EC), and Peru (PE). B) difference in monthly precipitation from climate normals at the three sites across the experimental periods for those sites. Bars indicate median values, black points indicate mean values. C) Cumulative precipitation over the course of the experimental period. Dotted lines indicate average cumulative precipitation, shaded regions indicate 95%, 80% and 50% quantiles. Climate normals are based on monthly precipitation for a period from 1958 to 1998

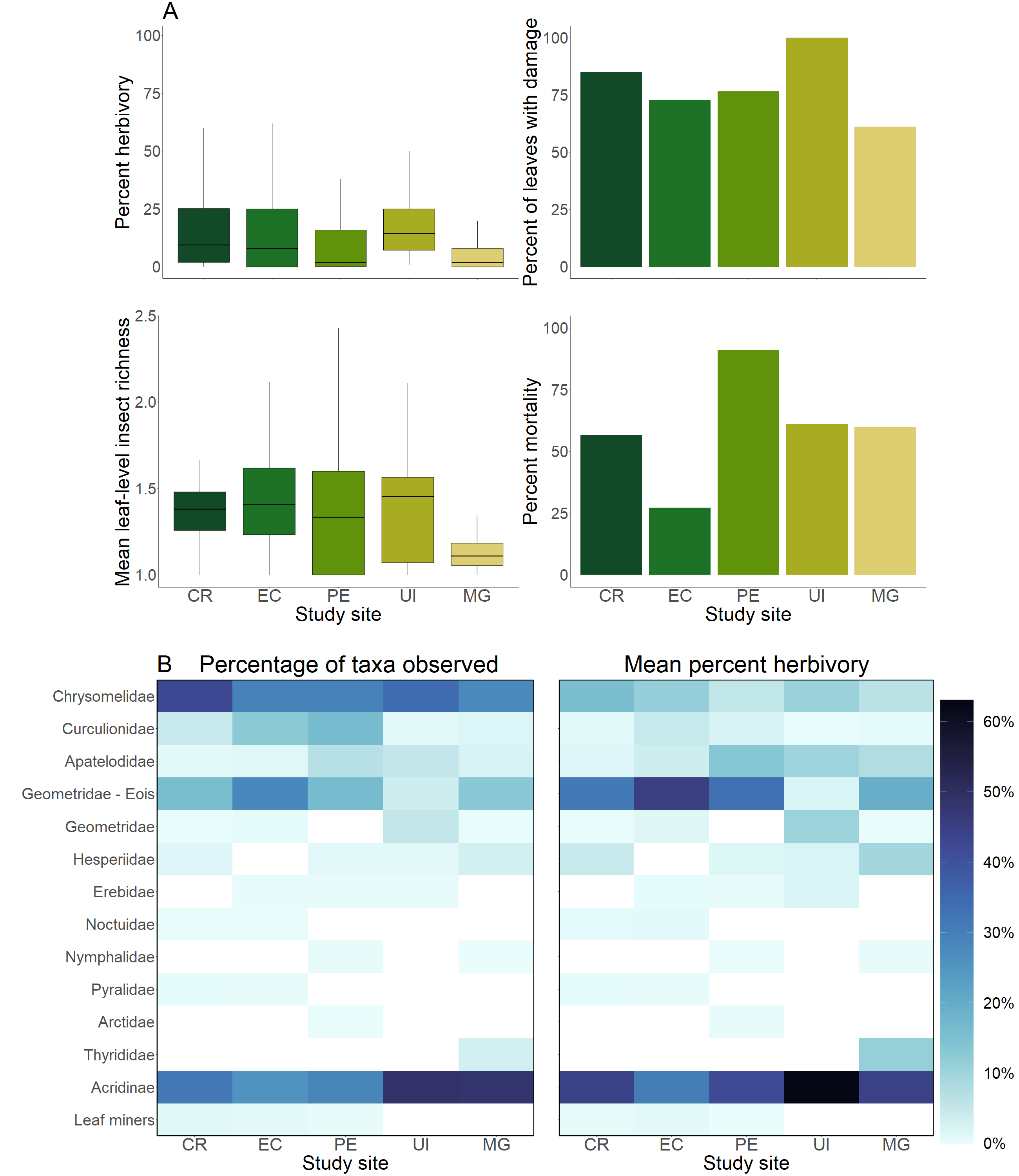

**Figure S3.** *Overall herbivory, plant mortality, and insect richness at five study sites*

A) Percent herbivory, percentage of leaves with any damage, herbivorous insect richness, and percent mortality of *Piper* in Costa Rica (CR), Ecuador (EC), Peru (PE), Uaimii (UI), and Mogi-Guaçu (MG), across treatments. B) Proportion of herbivorous insect taxa observed at each site measured by feeding damage patterns. Colored regions indicate the percentage of damage

observations contributed by each taxon. C) Proportion of herbivory attributed to each taxon at five study sites. Colored regions represent the percentage of each leaf consumed by each taxon at each site. White tiles represent sites where no leaf damage by that insect taxon was observed

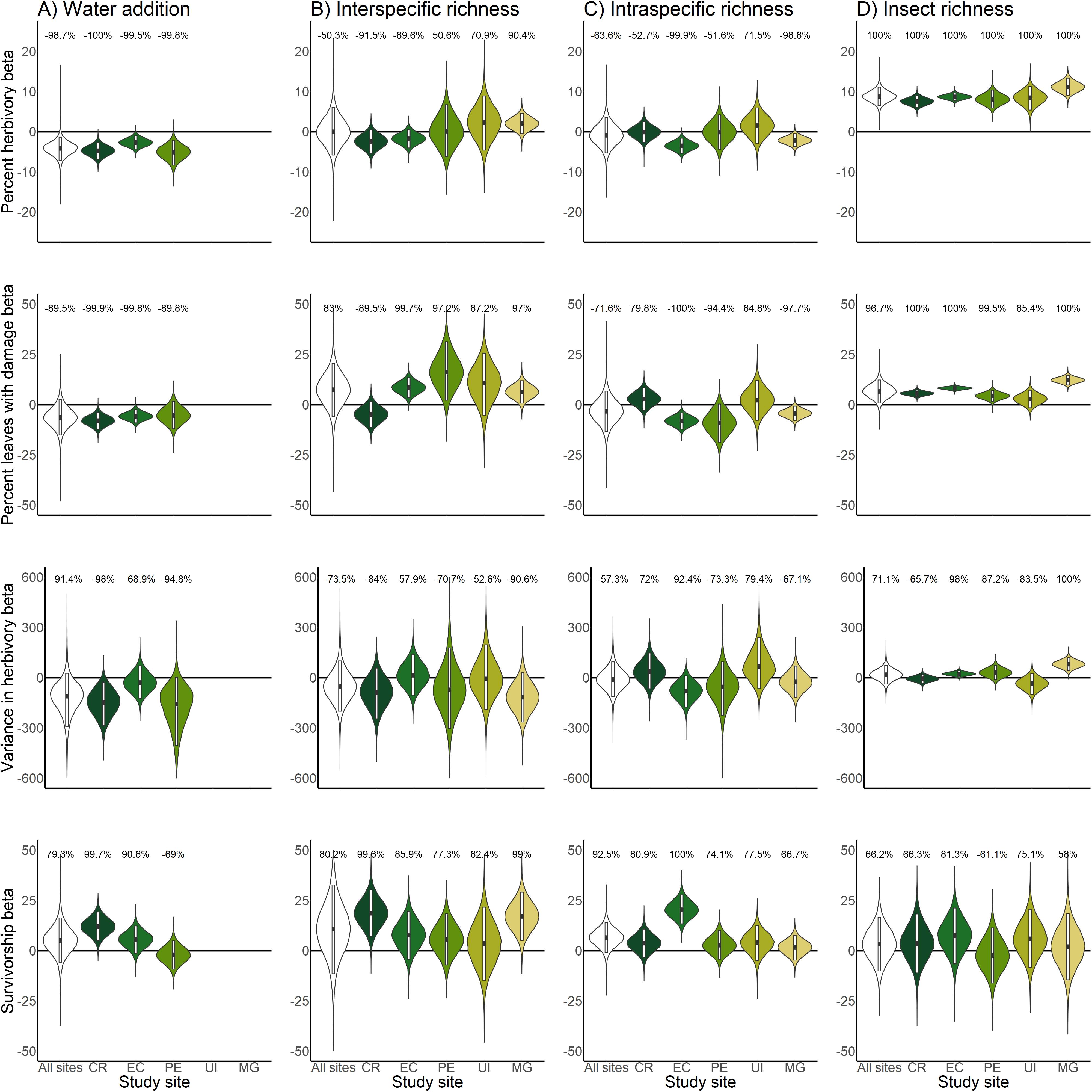
**Figure S4.** *HBM parameter estimates of percent herbivory, percentage of leaves with damage, variance in herbivory, and percent* Piper *survival against levels of A) water addition, B)* Piper *intraspecific richness, C)* Piper *interspecific richness, and D) insect richness at each site.*

Violins represent the posterior parameter distribution for each relationship across sites and within sites in Costa Rica (CR), Ecuador (EC), Peru (PE), Uaimii (UI), and Mogi-Guaçu (MG). Black lines represent the median posterior estimate while white bars represent 95% credibility intervals. Percentages above violins indicate the probability of an effect being entirely positive or entirely negative in response to an increase of the manipulated variable. Distributions for water addition compare unwatered control and watered plots; distributions for interspecific richness compare *Piper* species richness and are standardized as a percentage of the high richness treatment level; distributions for intraspecific richness compare low and high intraspecific richness plots; distributions for insect richness compare responses per insect taxon present at the leaf level

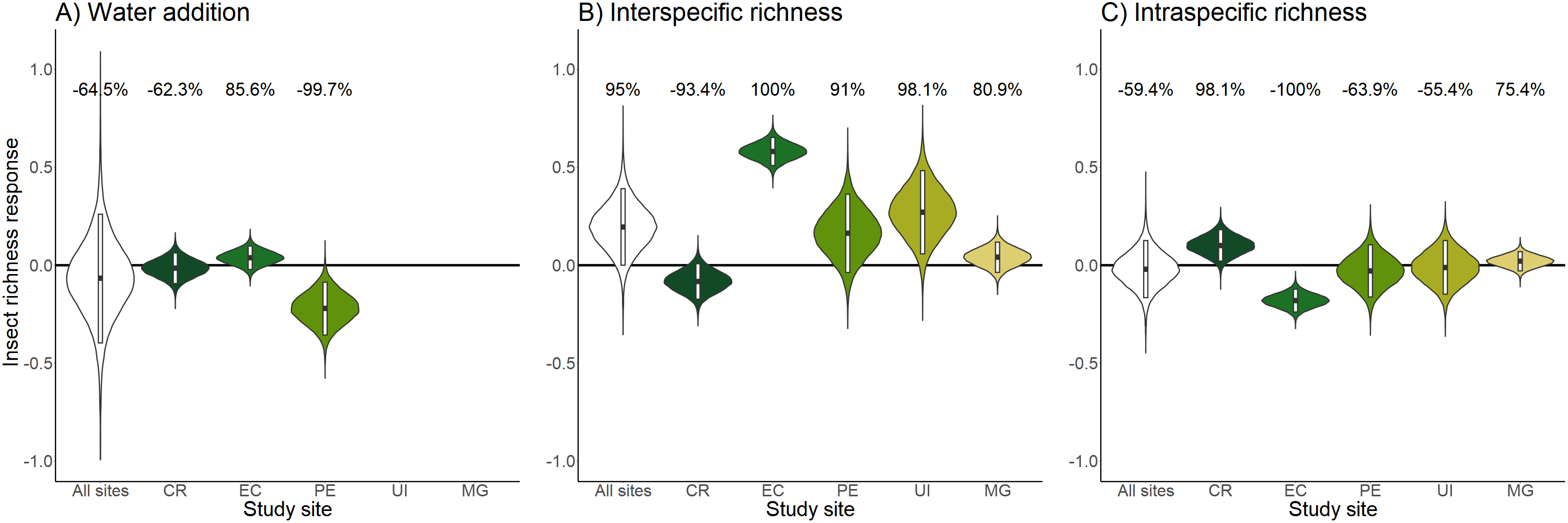
**Figure S5.** *HBM posterior parameter estimates of insect richness compared to levels of A) water addition, B)* Piper *intraspecific richness, and C)* Piper *interspecific richness*

Violins represent the posterior parameter distribution for each relationship across sites and within sites at Costa Rica (CR), Ecuador (EC), Peru (PE), Uaimii (UI), and Mogi-Guaçu (MG). Black lines represent the median posterior estimate while white bars represent 95% credibility intervals. Percentages above violins indicate the probability of an effect being entirely positive or entirely negative in response to an increase of the manipulated variable. Distributions for water addition compare the unwatered control and watered plots; distributions for interspecific richness compare *Piper* species richness and are standardized as a percentage of the high richness treatment level; distributions for intraspecific richness compare low and high intraspecific richness plots

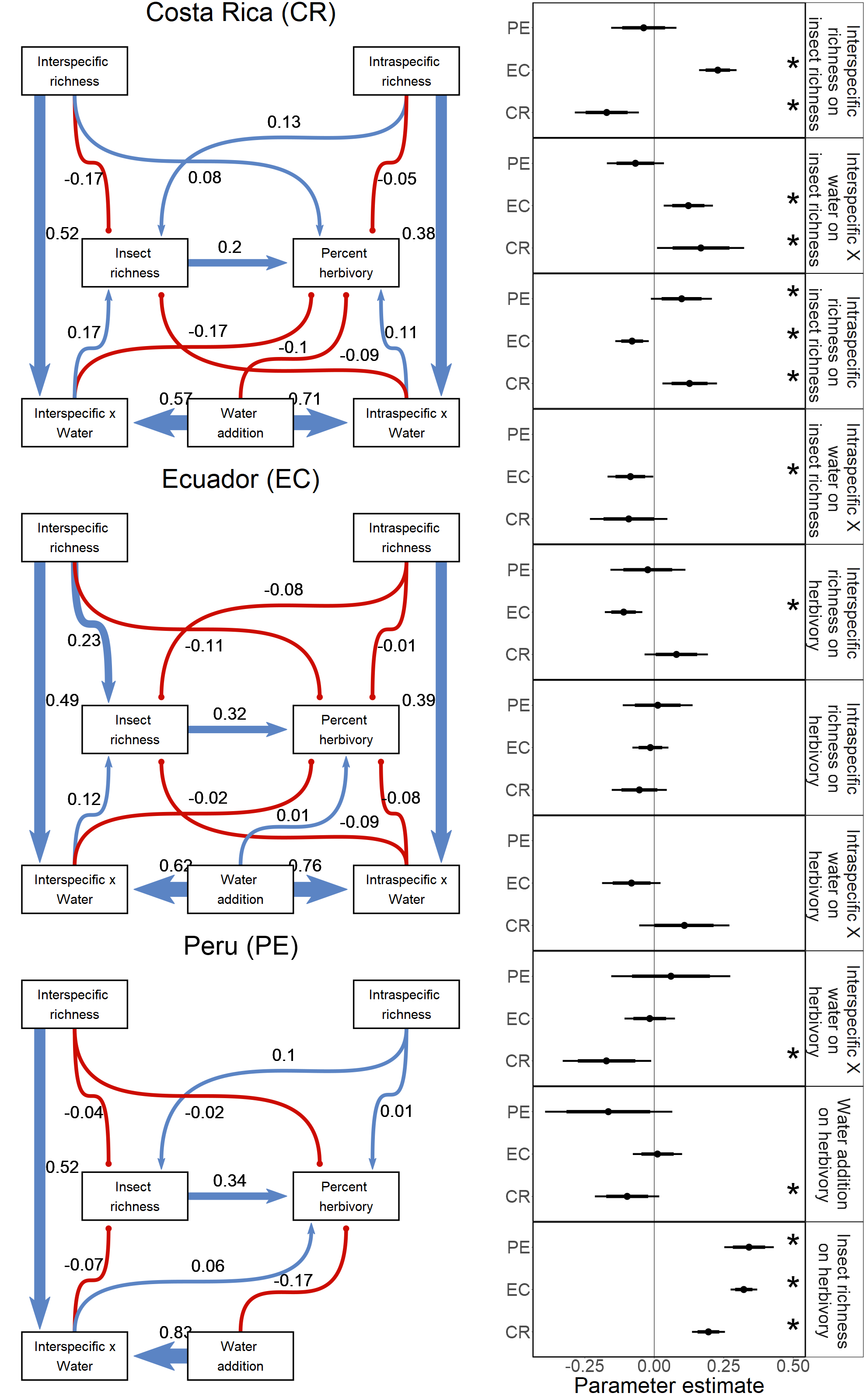

**Figure S6.** *Bayesian structural equation models for drivers of insect richness, herbivory and Piper survival at three sites, including interactions between intraspecific and interspecific richness, and water addition.* Path coefficients indicate the standardized mean of the posterior distribution for each causal path. Positive relationships are indicated in blue with triangular heads, and negative relationships are indicated in red with circular heads. Dot plots indicate the standardized mean of the posterior distribution for each causal path in A with 95% and 80% credibility intervals. Asterisks indicate causal paths where the probability of an effect being entirely positive or entirely negative is > 95%

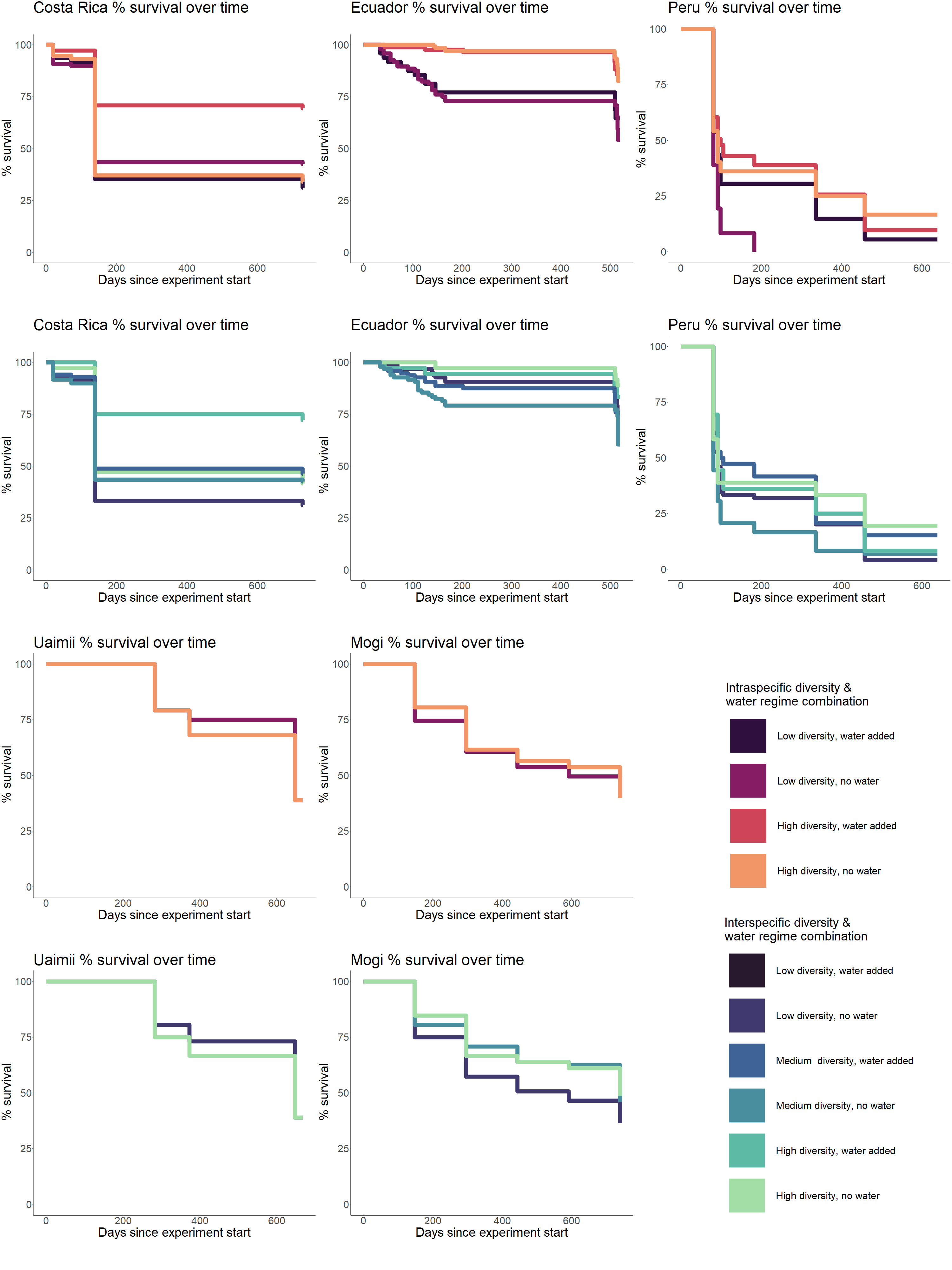
**Figure S7.** *Percent Piper survival over time in five sites, compared to levels of intraspecific richness and water addition*
